# Fluorescence Labeling of Circulating Tumor Cells with Folate Receptor Targeted Molecular Probes for Diffuse *In Vivo* Flow Cytometry

**DOI:** 10.1101/2020.01.22.913137

**Authors:** Roshani Patil, Madduri Srinivasarao, Mansoor Amiji, Philip S. Low, Mark Niedre

**Affiliations:** Department of Bioengineering, Northeastern University, Boston, MA, 02115, USA; Department of Chemistry, Purdue University, West Lafayette, IN, 47906, USA; Department of Pharmaceutical Sciences, School of Pharmacy, Northeastern University, Boston, MA, 02115, USA

**Keywords:** Circulating tumors cells (CTCs), folate receptor, diffuse imaging, fluorescence, *in-vivo* flow cytometry

## Abstract

**Purpose:** We recently developed a new instrument called ‘diffuse *in vivo* flow cytometry’ (DiFC) for enumeration of rare fluorescently-labeled circulating tumor cells (CTCs) in small animals without drawing blood samples. Until now, we have used cell lines that express fluorescent proteins, or were pre-labeled with a fluorescent dye *ex-vivo*. In this work, we investigated the use of two folate receptor (FR)-targeted fluorescence molecular probes for *in vivo* labeling of FR+ CTCs for DiFC.

**Methods:** We used EC-17 and Cy5-PEG-FR fluorescent probes. We studied the affinity of these probes for L1210A and KB cancer cells, both of which over-express FR. We tested the labeling specificity in cells in culture *in vitro*, in whole blood, and in mice *in vivo*. We also studied detectability of labeled cells with DiFC.

**Results:** Both EC-17 and Cy5-PEG-FR probes had high affinity for FR+ CTCs in cell culture *in vitro*. However, only EC-17 had sufficient specificity for CTCs in whole blood. EC-17 labeled CTCs were also readily detectable in circulation in mice with DiFC.

**Conclusions:** This work demonstrates the feasibility of labeling CTCs for DiFC with a cell surface receptor targeted probe, greatly expanding the utility of the method for pre-clinical animal models. Because DiFC uses diffuse light, this method could be also used to enumerate CTCs in larger animal models and potentially even in humans.

## 1. Introduction

Cancer metastasis is a multi-step process, by which tumor cells colonize distant organs and tissues. The circulatory system is one of the most common pathways, wherein tumor cells intravasate into the peripheral blood, circulate, and form metastases at secondary sites [1–3]. Circulating tumor cells (CTCs) in the bloodstream are therefore of great interest in cancer research. Numerous studies have demonstrated that CTC numbers correlate with overall survival, disease progression, and response to treatment for many cancers [4–9]. CTCs are extremely rare; fewer than 1 CTC per mL of peripheral blood is considered prognostically negative [10].

Normally, CTCs are studied using ‘liquid biopsy’, where relatively small blood samples are drawn from the patient or small animal (in the case of pre-clinical research), are purified, enriched and further analyzed [11–12]. Although these are widely used in biomedical research, they are far from optimal for a number of reasons. In particular, small blood samples provide poor statistical sampling of the circulating blood volume [13–15], blood is known to degrade rapidly after removal from the body [16], and enrichment can cause cell loss or dissolution [17–18]. Moreover, in the case of small animal studies CTCs may be so rare that it is necessary to draw and analyze the entire blood volume, which requires euthanizing the animal [19]. This precludes longitudinal study of individual animals over time.

These limitations have driven the development of optical methods for enumerating circulating cells without having to draw blood samples, collectively termed ‘*in vivo* flow cytometry’ (IVFC) [20–23]. IVFC typically uses specialized confocal microscopy [24–27] or photoacoustic [28–30] instrumentation to detect circulating cells, for example in a small blood vessel in the ear of a mouse. Our group recently developed ‘diffuse *in vivo* flow cytometry” (DiFC) [31–34], a new technique for counting rare fluorescently-labeled CTCs with highly scattered light. This allows sampling of large circulating blood volumes (compared to a microscope), and therefore detection of less abundant cells. For example, we recently [33] used DiFC to monitor dissemination of multiple myeloma cells in a xenograft model, and showed that we could non-invasively detect fewer than 1 CTC per mL of blood. In addition to detection sensitivity, a major advantage of DiFC is that it works in bulk, optically diffusive tissue such as the mouse leg or tail (as opposed to the thin ear of a mouse as in microscopy) so in principle could be used in larger limbs and species. However, until now we have only used DiFC with CTCs that express green fluorescent proteins (GFP) or by labeling cells *ex-vivo* with membrane or cytosol dyes. This presently limits the use of DiFC to the study of mouse xenograft models using cultured immortalized cell lines [32–33].

Use of a targeted fluorescent molecular probe that could label CTCs while in circulation would therefore greatly expand the utility of DiFC. In this work, we studied two folate receptor (FR) alpha (α) targeted molecular probes for DiFC for the first time. FR is widely used as a therapeutic and diagnostic target for cancer, since it is often over-expressed in many epithelial cancers [35–38], including ovarian [39–42], breast [43–44], and non-small cell lung carcinomas [45–46]. It also is purported to have very low expression in normal tissues [35–37]. Cell surface-receptor targeted molecular contrast agents have already been used for microscopy-IVFC previously [22, 47] including for FR+ cells [24, 48]. However, a major question was whether CTCs could be labeled *in vivo* with sufficient specificity and brightness for detection with DiFC. DiFC uses diffuse light and is more therefore susceptible to scatter and attenuation of biological tissue (compared to microscopy) [49], and as such may have higher fluorescence labeling requirements [22, 32].

Specifically, we used EC-17 and Cy5-PEG-FR FRα-fluorescent probes. EC-17 is a small-molecule FITC based probe that was originally developed for fluorescence guided surgery (FGS) and has been shown to have high-affinity for FR+ CTCs previously [48]. Moreover, EC-17 and its NIR analog OTL-38 have both been used in FGS clinical trials [39, 50], leading to the exciting possibility of using DiFC in humans in the future. Cy5-PEG-FR is a larger molecular weight probe, but has the advantage of having red excitation and emission wavelengths, which in principle is favorable for DiFC because of reduced optical attenuation of red light in biological tissue [49]. We tested these probes with two FR+ human cancer cell lines. As we show, EC-17 has very sensitivity and specificity for FR+ CTCs and labeled CTCs were detectable in mice *in vivo* with DiFC. In contrast, Cy5-PEG-FR demonstrated significant non-specific uptake in blood. Overall, this work demonstrates the feasibility of using targeted molecular probes for DiFC for the first time, which could greatly extend its utility in CTC research.

## 2. Materials and Methods

### 2.1 Diffuse In Vivo Flow Cytometry

The DiFC instrument setup is shown in ***figs. 1a, b***. The design and signal processing algorithms were described in detail by us previously [32–33]. Briefly, DiFC works on the principle of laser-induced fluorescence. The DiFC fiber probes (red arrows, ***fig. 1c,d***) have integrated optical filters and lenses that allow detection of the weak fluorescence signal from individual moving cells in the bloodstream [34]. When placed over a major blood vessel, for example in the tail or leg (***fig. 1c***) of a mouse, transient fluorescence peaks are detected and counted We showed previously that the ventral caudal artery in a mouse tail carried hundreds of μL of blood per minute [32], allowing us to detect and count very rare circulating cells [33].

**Figure 1:**
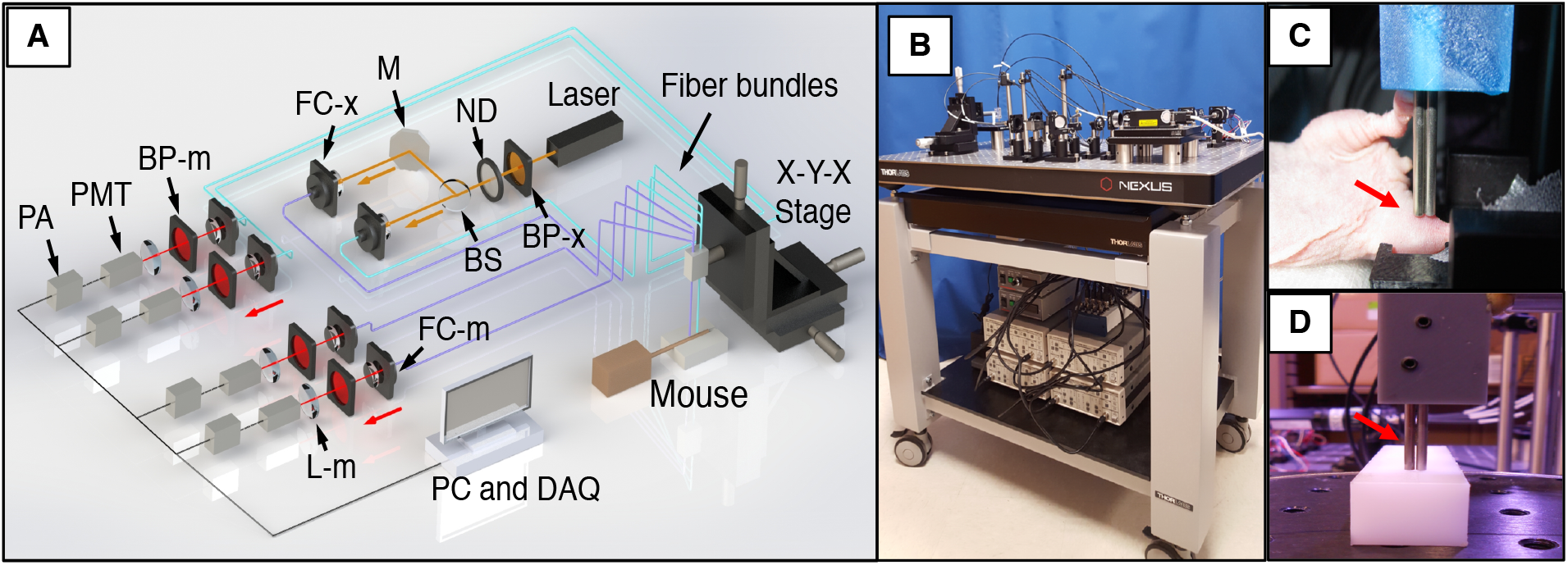
(a) DiFC instrument schematic, reproduced with permission from X. Tan et. al. [32]. See text for details. (b) photograph of the b-DiFC cart mounted system, reproduced with permission from R. Patil et. al. [33]. The dual-fiber DiFC probe head (red arrows) is placed on the surface of the sample, in this case either (c) the hindleg of a mouse (femoral artery), or (d) a tissue mimicking flow phantom. This permits detection and enumeration of fluorescently-labeled circulating cells.

We previously built blue (b-DiFC) and red (r-DiFC) versions of DiFC that are compatible with absorption and emission spectra of FITC and Cy5.5 fluorescent dyes used in this study (see ***section 2.2***). A number of minor hardware improvements were made to the b-DiFC system here. First, we used voltage-output photomultiplier tubes (Hamamatsu H10722-20; Edmund Optics, Barrington, NJ) and voltage pre-amplifiers (SR650; Stanford Research Systems, Sunnyvale, CA) rather than current-output PMTs, due to improved noise properties. Second, we added a second, 536 nm bandpass interference filter (IDEX Health and Science, LLC, Rochester, NY) in front of each PMT to improve out-of-band blocking.

### 2.2 Folate-Receptor (FR) Targeted Fluorescent Molecular Probes

EC-17 is a small molecule FRα targeted probe (MW: 917 Da) which was developed in the lab of Prof. Low, and been extensively characterized previously [48, 51]. EC-17 is a conjugation of Fluorescein Isocyanate (FITC) to folic acid, which is the binding ligand of folate receptor. Previous work showed that EC-17 has excellent affinity for FR+ CTCs in blood compared to larger antibody-based probes [24, 48]. The EC-17 maximum excitation and emission wavelengths are 490 nm and 520 nm, respectively.

Cy5-PEG-FR is a larger (MW: 3500 Da) FRα-targeted probe (PG2-FAS5-2k; Nanocs Inc, New York, N). Cy5-PEG-FR has a 2 kDa PEG linker chain between the Cy5 and folate groups, which maintains its photostability and avoids Cy5 quenching, however as we show may contribute to non-specific uptake in blood. The maximum excitation and emission wavelengths of Cy5-PEG-FR are 647nm and 664nm, respectively.

### 2.3 Cell lines

KB are FR+ HeLa-derived human cervical cancer line and were purchased from ATCC (CCL-17; ATCC, Manassas, VA). L1210A is an FR+ human leukemia cancer line that was previously modified to express folate receptors (Purdue University, West Lafayette, IN). MM.1S multiple myeloma cells were used as an FR-control and were also purchased from ATCC (CRL-2974). All cell lines were cultured in RPMI 1640, folate deficient media (Gibco 27016-021; ThermoFisher Scientific, Waltham, MA).

### 2.4 Fluorescent Microspheres

We used fluorescence microspheres as a reference standard for evaluating the brightness of labeled cells. “Dragon Green” (DG; DG06M, Bangs Laboratories, Fishers, IN) and “Flash Red” (FR; FR06M, Bangs) microspheres are sold in kits of 5 intensities, ranging from 1 (lowest) to 5 (highest). These serve as useful standards for evaluating cell labeling, and for comparing data between different instruments, for example between a flow cytometer (FC) and DiFC. Our previous work showed that cells with fluorescent labeling equal to or exceeding FR4 had high signal-to-noise ratios *in vivo* [32]. Likewise, blue fluorescence labeling exceeding DG3 were readily detectable with b-DiFC in SCID mice *in vivo* [33].

### 2.5 Labeling of Cells with FR-Targeted Probes *In Vitro*

We first tested labeling of FR+ CTC with the FR-targeted probes in cells in culture *in vitro*. EC-17 was added at a concentration of 200 nM to suspensions of 10^6^ cells/mL in 2% FBS in PBS in a 6-well plates. Cells were incubated at 37°C for 60 minutes, then washed twice with PBS, and resuspended at a concentration of 10^6^ cells/mL. Cy5-PEG-FR was added at a concentration of 1.6 μM to suspensions of 0.5 × 10^6^ cells/mL at 4°C for 30 minutes. Following this, cells were washed twice and resuspended at a concentration of 10^6^ cells/mL prior to testing with flow cytometry (FC) or DiFC.

### 2.6 Labeling of Cells in Whole Mouse Blood

We next tested labeling of FR+ CTCs in whole mouse blood. Blood was drawn from female nude (nu/nu) mice (Charles River Laboratory, Wilmington, MA), which were kept on folic-acid free diet (TD.00434; Teklad Diet, UK) for at least 2 weeks prior to the study to minimize free folic acid in the blood. Drawn blood was stabilized and diluted 1:1 with 1000 units/ml Heparin (Sigma Aldrich, Natick, MA). 10^4^ CTCs were added to 500 μL blood aliquots (“spiked”) and then gently agitated for 30 seconds to ensure that samples were well mixed. 200 nM EC-17 or 1.6 μM Cy5-PEG-FR were added to the blood samples, and these were incubated for 60 minutes as above.

To test EC-17 labeling specificity, target (KB, L1210A or MM.1s) cells were first pre-labeled with Cell Trace Far Red (CTFR) dye (C34564, Molecular Probes Inc., Eugene, OR) according to the manufacturer’s instructions. The rationale here was that well EC17-labeled cells should exhibit double (blue and red) positive fluorescence on FC. Likewise, to test the specificity of Cy5-PEG-FR labeling, target cells were pre-labeled with Cell Trace CSFE dye (C34570, Molecular Probes) according to the manufacturer’s instructions.

Some samples were also co-incubated with 10 μM of free folic acid to introduce competitive binding. Blood suspensions were then analyzed using the blue (488 nm laser, 530 nm emission filter), and red (637 nm laser, 647 nm emission filter) channels of an Attune NXT flow cytometer (ThermoFisher). Experiments were repeated at least in duplicate in each case.

### 2.7 DiFC Flow Phantom Experiments *In Vitro*

As an initial test of the detectability of EC-17 and Cy5-PEG-FR labeled cells with DiFC, we used a tissue-simulating flow phantom model (***fig. 1d***), as we have in our previous work [32]. Briefly, the flow phantom was a block of optically diffusing high-density polyethylene (HDPE) with similar scatter and absorption properties of biological tissue. A strand of microbore Tygon tubing (TGY-010-C, Small Parts, Inc., Seattle, Washington) was embedded in the phantom block at a depth of 0.75 mm. The tubing was connected to a syringe pump (70-2209, Harvard Apparatus, Holliston, Massachusetts). We prepared suspensions of 10^3^ per mL cells suspension of EC-17 and Cy5-PEG-FR labeled L1210A cells per mL, labeled as described in section 2.4. Cell suspensions were run through the phantom at a rate of 50 μL/s. We also tested 10^3^ spheres/mL suspensions of DG3 and FR4 fluorescence reference microspheres at concentrations of 10^3^ spheres/mL as a comparison.

### 2.8 DiFC Mouse Experiments *In Vivo*

All mice were handled in accordance with Northeastern University’s Institutional Animal Care and Use Committee (IACUC) policies on animal care. Animal experiments were carried out under Northeastern University IACUC protocol #15-0728R. All experiments and methods were performed with approval from, and in accordance with relevant guidelines and regulations of Northeastern University IACUC.

L1210A leukemia cells were used in these experiments, since they are known to circulate for extended periods of time when intravenously injected in mice [14]. We first labeled L1210A cells with EC-17 in vitro as above. 10^6^ labeled cells were suspended in an injection volume of 200 μL of cell culture media. Female nude mice were anesthetized with isofluorane, and 10^6^ cells the cell suspension was injected *i.v.* via the tail vein (N = 3). We performed DiFC on the femoral blood vessel in the mouse leg (***fig. 1e***), beginning 10 minutes after injection of the cells. We used a detection threshold of 6 times the standard deviation of the background signal, which was calculated for each minute of the trace. As we discuss this virtually eliminated detection of false positive signals due to electronic noise or motion artifacts. At the end of the DiFC scan, approximately 1 mL of blood was harvested (terminal) and EC-17+ cells counted by flow cytometry.

Second, we also performed proof-of-principle testing of EC-17 labeling of FR+ CTCs while in circulation (which we subsequently refer to as “*in vivo* labeling”). Specifically we injected 10^6^ L1210A cells *i.v.* via the tail vein. 5 minutes later, we administered 25ug of EC-17 probe by *i.v*. injection. We performed DiFC on the mouse hindleg, beginning approximately 40 minutes after the probe injection (45 minutes after cell injection).

## 3 Results

### 3.1 Labeling of FR+ CTCs in Cells *In Vitro*

We first tested labeling of FR+ CTCs with EC-17 in cell culture *in vitro*. ***Figure 2a*** shows that FR+ CTCs (L1210A and KB) cells were brightly labeled, with 94.7% of EC-17-labeled L1210A cells and 94.1 % of labeled KB cells exceeding the fluorescence intensity of Dragon Green 3 (DG3) microspheres. As noted above, DG3 are a reference standard that we often use to compare the brightness of labeled cells across instruments; we have shown previously that cells with brightness exceeding DG3 are readily detectable *in vivo* with DiFC. In contrast, FR− MM.1s cells displayed negligible uptake, with comparable fluorescence to unlabeled L1201A and KB cells, illustrating the specificity of the EC-17 probe.

**Figure 2:**
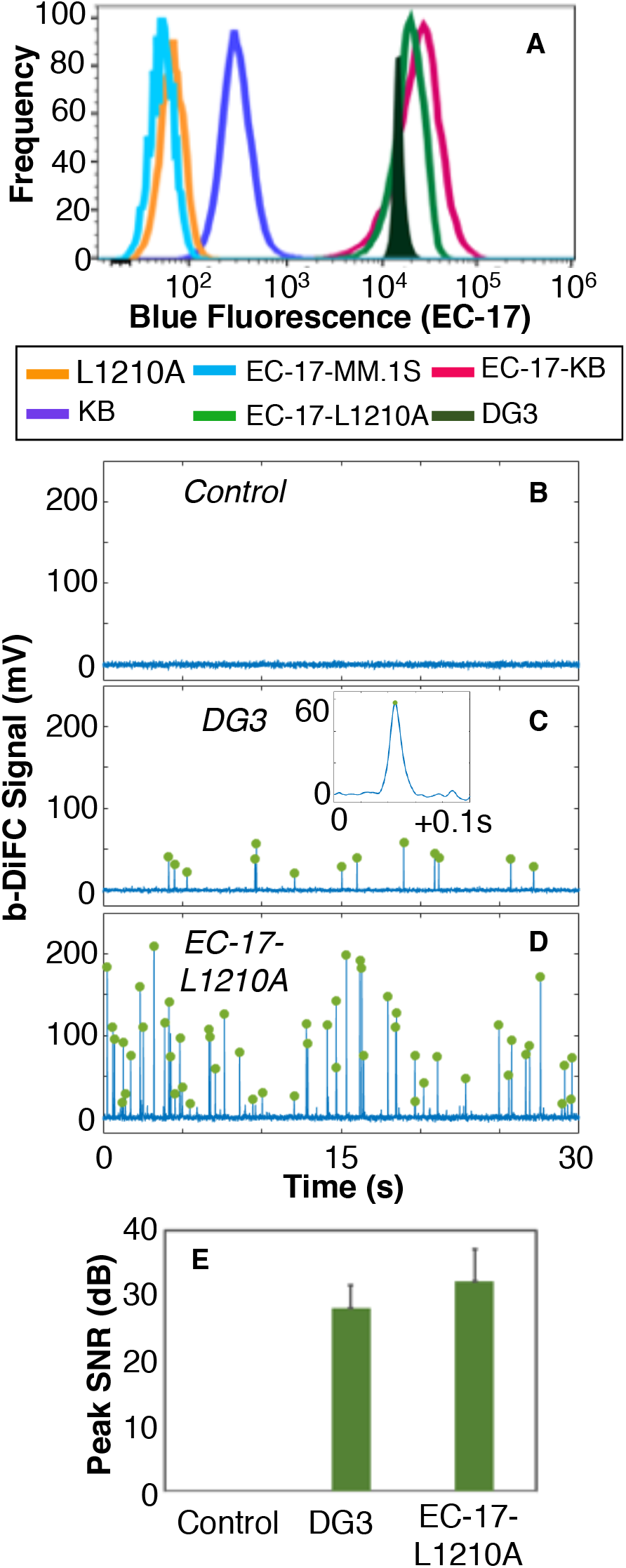
[single column figure] (a) blue-channel fluorescence flow cytometry (FC) analysis of EC-17 labeling of cells in culture in vitro are shown. EC-17 had high affinity for FR+ L1210A and KB cells, whereas little uptake by FR− MM.1s cells was observed. Representative b-DiFC measurements from (b) unlabeled (control) L1210A cells in culture in a flow phantom, (c) DG3 reference fluorescent microspheres, and (d) EC-17 labeled L1210A cells, which were readily detectable with b-DiFC. (e) Summary of the mean peak amplitudes measured with b-DiFC in each case are shown.

We next tested if EC-17-labeled FR+ CTCs were sufficiently brightly-labeled to be detectable with b-DiFC in a tissue-simulating optical flow phantom (***fig. 1d***) [33]. Representative b-DiFC scans of un-labeled L1210A cells (controls), DG3 microspheres, and EC-17-labeled L1210A cells, are shown in ***figs 2b-d***, respectively. Each fluorescent peak in ***figs. 2c,d*** corresponds to the detection of a fluorescently labeled cell passing through the DiFC field of view. The inset panel in ***fig. 2c*** shows the width of a representative peak. As summarized in ***fig. 2e***, the b-DiFC-measured amplitude of EC-17 labeled cells was on average greater than DG3 microspheres, which was consistent with the FC fluorescence data. Here, the signal to noise ratio (SNR in dB) is defined as ***SNR = 20log10(I/σ)***, where ***I*** is the mean peak amplitude (intensity), and ***σ*** is the DiFC instrument noise.

We next tested labeling of L1210A and MM.1s cells with Cy5-PEG-FR, as shown in ***fig. 3a***. The fluorescence intensity of FR4 reference microspheres (which, as above, approximates a well-labeled cell that produces a high SNR when detected with DiFC *in vivo*) is also shown for comparison. By inspection, there was a large range of cell labeling observed with Cy5-PEG-FR, with 21.7% of cells exceeding the brightness of FR4. We also performed r-DiFC for un-labeled (control) L1210A cells, FR4 microspheres, and Cy5-PEG-FR-labeled L1210A cells as shown in ***figs. 3b-d***, respectively. Although Cy5-PEG-FR-labeled L1210A cells were clearly detectable in the phantom, there was a broad range of detected peak intensities with r-DiFC, which was (again) consistent with the FC data. The mean measured r-DiFC peak SNRs are summarized in ***fig. 3e***.

**Figure 3:**
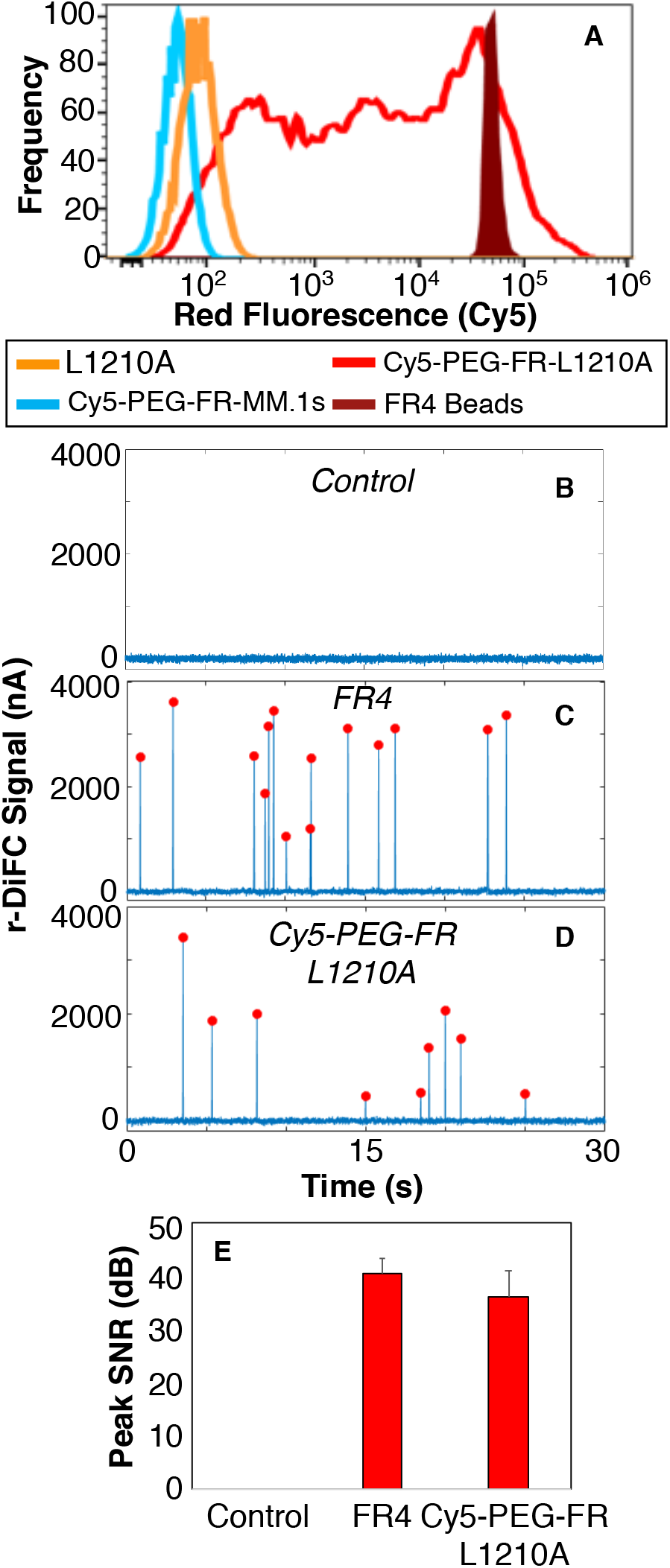
[single column figure] (a) red-channel fluorescence flow cytometry (FC) analysis of Cy5-PEG-FR labeling of cells in culture in vitro are shown. Cy5-PEG-FR had reasonable affinity for FR+ L1210A cells, although a wide range of labeling efficiency was shown. Likewise, little uptake by FR− MM.1s cells was observed. Representative r-DiFC measurements from (b) unlabeled (control) L1210A cells in culture in a flow phantom, (c) FR4 reference fluorescent microspheres, and (d) Cy5-PEG-FR labeled L1210A cells, which were also readily detectable with r-DiFC. (e) Summary of the mean peak amplitudes measured with r-DiFC in each case are shown.

### 3.2 Labeling of FR+ Cells in Whole Mouse Blood

We next tested labeling of FR+ CTCs in whole mouse blood with EC-17 and Cy5-PEG-FR probes. Blood is a complex suspension of billions of cells per mL, and therefore presents a more realistic model of in vivo labeling where non-specific cell uptake and competitive binding may occur.

The results are summarized in ***figure 4***. The blue (horizontal axis) and red (vertical axis) autofluorescence of whole mouse blood is shown in ***fig. 4a***. Addition of EC-17 probe to the blood (only) showed a small amount non-specific uptake of EC-17 (***fig. 4b***, ***blue*** horizontal axis), most likely by macrophages which is consistent with previously reported work [48]. As shown in ***fig. 4c***, we added CTFR-prelabeled L1210A cells to blood, and then EC-17 probe. More than 90% of L1210A CTCs were well labeled with EC-17, and the average brightness of labeled cells was above non-specific background levels (shown in ***fig. 4b)***. As shown in ***fig. 4d*** FR+ KB cells were also brightly labeled with EC-17. Furthermore, addition of free folic acid to blood samples spiked with L1210A cells (but prior to addition of EC-17) resulted in competitive blockage of subsequent probe binding (***fig. 4e***). In addition, FR− MM.1s cells showed little binding of the EC-17 probe (***fig 4f***).

**Figure 4:**
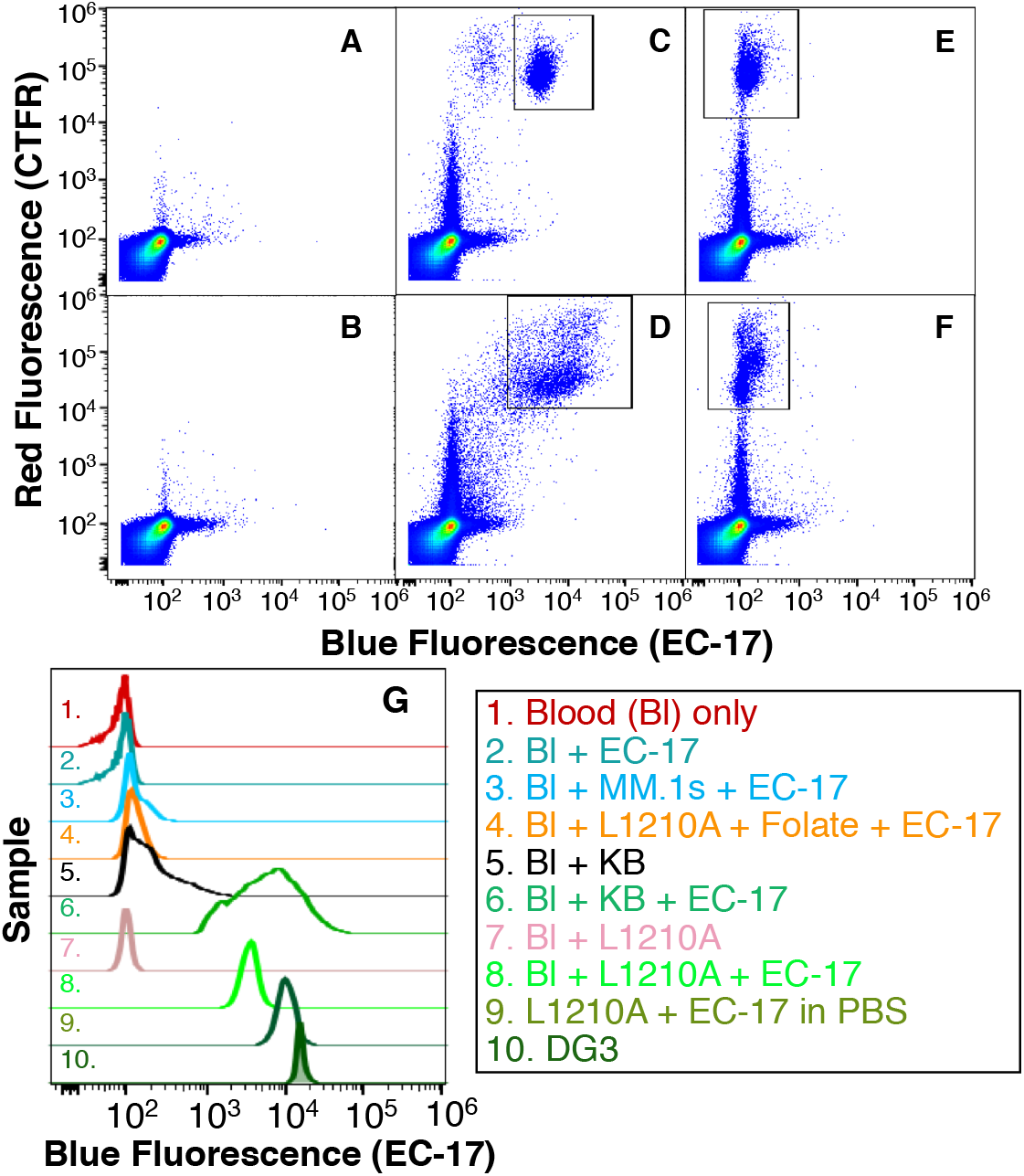
Labeling of CTCs with EC-17 in whole mouse blood in vitro. (a) Red and blue autofluorescence of blood (only). (b) Non-specific uptake of EC-17 by blood cells was minimal, whereas there was substantial specific labeling of FR+ (c) L1210A and (d) KB cells. (e) addition of free-folic acid blocked binding of EC-17 to L1210 cells in blood. (f) FR− MM.1s cells showed negligible EC-17 uptake. (g) Summary of blue-fluorescence FC measurements for all experiments performed is shown.

The distributions of EC-17 (blue) fluorescence for all the experiments performed in this study are summarized in ***fig. 4g***. When considering blue-fluorescence alone, these data show a clear separation between non-specific uptake of blood cells and FR− CTCs with EC-17 (lines 2 and 3) compared to FR+ CTCs labeling by EC-17 (lines 8 and 9). In addition, the maximum fluorescence labeling of CTCs in the blood was 2.9 times lower than labeling in simple PBS solution *in vitro* (***fig. 2g***), which is unsurprising given the relative complexity of blood versus PBS.

Using the same methodology, we tested Cy5-PEG-FR labeling of FR+ L1210A cells as summarized in ***figure 5***. Blood autofluorescence is shown in ***fig. 5a***. Addition of the Cy5-PEG-FR probe to whole blood showed significant non-specific uptake as shown in ***fig. 5b*** (vertical axis, red channel). Cy5-PEG-FR probe labeling of blood samples spiked with CFSE-labeled L1210A cells yielded a significant double-labeled population (***fig 5c***). However, the large non-specific uptake by blood cells (***figs. 5b,c***) in the same fluorescence intensity ranges meant that the target cell population would not be identifiable using only the red (Cy5) fluorescence as shown in ***fig. 5d***, and therefore would be unsuitable for r-DiFC.

**Figure 5:**
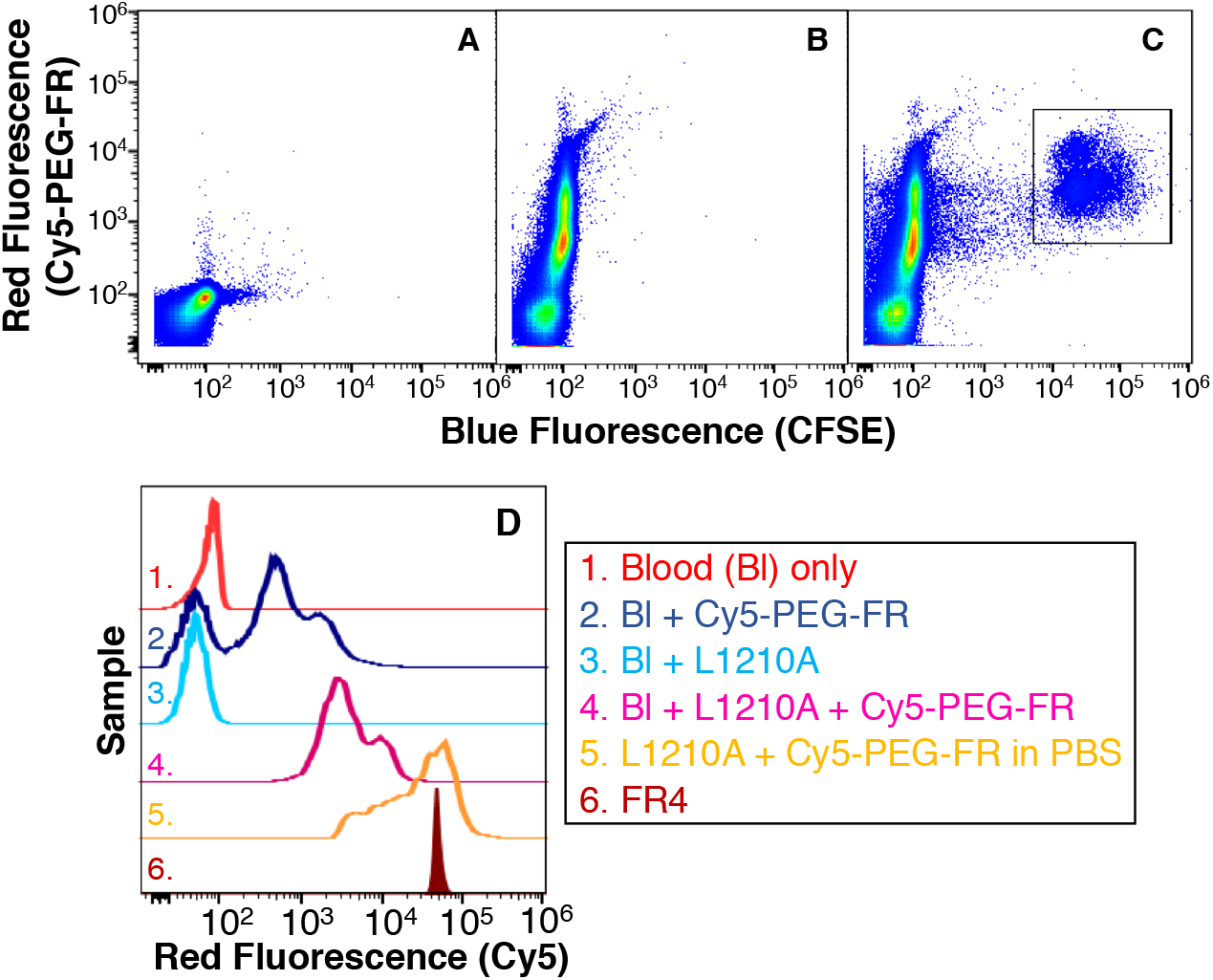
Labeling of CTCs with Cy5-PEG-FR in whole mouse blood in vitro. (a) Red and blue autofluorescence of blood (only). (b) Non-specific uptake of Cy5-PEG-FR by blood cells was significant.(c) Labeling of L1210A FR+ CTCs with Cy5-PEG-FR was significant, but (d) there was insufficient separation of FR+ labeling and non-specific uptake using red fluorescence alone.

In summary, these *in vitro* blood labeling experiments demonstrated that the small-molecule EC-17 was the more promising probe for *in vivo* labeling of FR+ CTCs for DiFC.

### 3.3 *In Vivo* Detection of EC-17 labeled CTCs with DiFC

We next tested if FR+ CTCs that were well-labeled with EC-17 were detectable with DiFC in mice *in vivo*. To test this, we pre-labeled L1210A cells with EC-17 in culture *in vitro*, and then injected these *i.v.* via the tail vein into nude mice. We performed b-DiFC on the mouse hind-leg, approximately above the large femoral blood vessel as in ***fig. 1c***. Representative b-DiFC data is shown in ***figure 6***. ***Fig. 6a*** shows b-DiFC from a non-injected control mouse. As shown, the background signal was quite stable, with a mean noise standard deviation (σ) of 1.9 mV, and false-alarm rate (FAR) of 0.008 counts per minute, when a detection threshold of 6 times σ was used (calculated for each mouse). ***Fig. 6b*** shows representative b-DiFC data from a mouse injected with pre-labeled L1210A cells. Labeled L1210A cells were clearly detectable with b-DiFC, with mean signal to noise ratio of 18 dB, although some individual detections exceeded 30 dB. The mean count rate for all 3 mice tested was 0.8 cell counts per minute (48 counts per hour).

**Figure 6:**
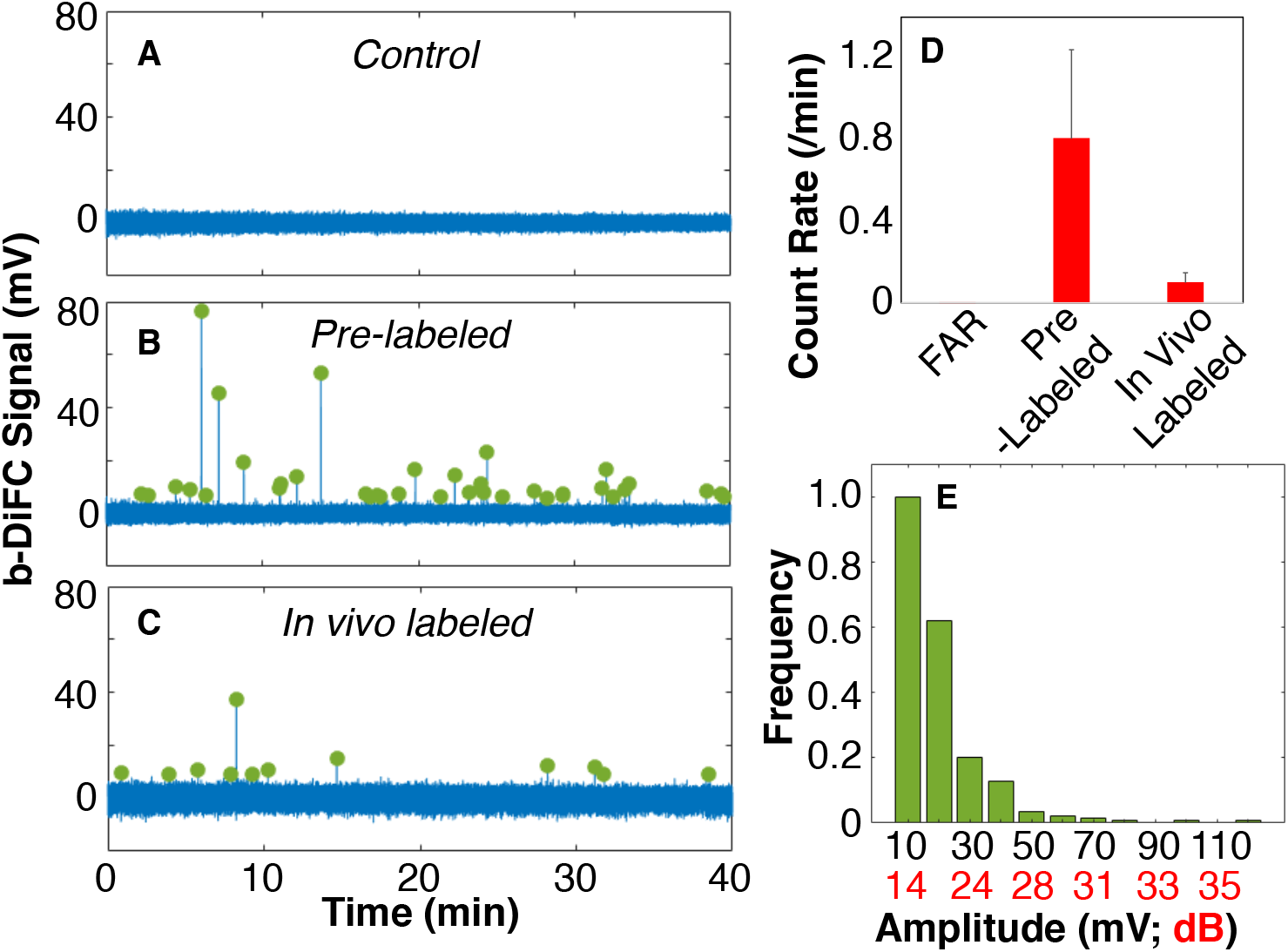
(a) Representative b-DiFC measurement from the hindleg of an un-injected control mouse. (b) L1210A cells that were pre-labeled in culture with EC-17 were readily detectable in mice in vivo. (c) L1210A cells and EC-17 were injected 5 minutes apart so that cells were labeled in vivo. (d) Summary of the FAR and in vivo b-DiFC count rates for N=3 mice tested in each case. (e) Histogram of peak amplitudes for EC-17 labeled cells.

Following DiFC scanning, we drew 1 mL of blood (terminal) from the mice and counted EC-17+ cells in the blood using fluorescence (blue channel) FC. On average we counted 13.2 ± 5.7 EC-17 labeled cells per mL in blood, where we defined “labeled” as exceeding the blue fluorescence intensity of DG3 microspheres. This relatively low number of cells in the blood was expected from the fact that blood was sampled approximately 100 minutes after injection of the cells, meaning that very few L1210A cells remained in circulation.

Using this cell concentration, we were able to estimate the DiFC blood sampling rate from the mouse leg (femoral artery) as follows: The DiFC count rate measured during the final 10 minutes of the scan (just before drawing blood) was 0.6 ± 0.3 counts per minute. Compared to the estimated number of L1210A cells in the blood (13.2 cells/mL), this implies that DiFC sampled approximately 0.6 counts min^−1^/ 13.2 cells mL^−1^ = 48 μL / minute of blood. This is lower than our previously reported flow speed in ventral caudal artery in the mouse tail [32], which reflects the smaller size of the mouse femoral artery compared to the tail artery.

### 3.4 *In Vivo* Labeling of CTCs with EC-17 and Detection with DiFC

Direct *in vivo* labeling of FR+ CTCs while in is a more challenging problem, and based on our studies in mouse blood samples we anticipated that CTC labeling would be significantly less bright than in culture *in vitro*. To test this, we first injected a suspension of L1210A cells in mice. 5 minutes later, we injected EC-17 probe. After approximately 40 minutes (allowing time for the free EC-17 to clear from circulation), we performed b-DiFC on the mouse hind-leg as above. Representative data is shown in ***fig. 6c***. As shown, we were able to detect a small number of cells in circulation, at an average count rate of 0.1 per minute.

The measured FAR and count rates for pre-labeled and in-vivo labeled cells are summarized in ***fig. 6d***. The lower count rate for in vivo labeling (12.5% of pre-labeled) stems from two reasons. First we started DiFC scanning approximately 45 minutes after injection of cells (versus 10 minutes for pre-labeled cells), so that many of the injected cells likely already cleared from circulation. Second, uptake and labeling of EC-17 cells by L1210A cells while in circulation *in vivo* was likely significantly less efficient than solution.

With respect to the latter, the histogram of detected peak amplitudes (expressed in mV and dB) *in vivo* are summarized in ***fig. 6e***. From these data, we can estimate (approximately) that the lower count rate for *in vivo* labeled cells could be explained by a reduction of 14-16 dB labeling brightness. This in turn suggests that cells labeled *in vivo* were approximately 5-to-6 times lower in fluorescence brightness (labeling) than cells that were pre-labeled in culture. This is a reasonable estimation, given that we observed a reduction in EC-17 labeling brightness by a factor of 2.9 when labeling in blood compared to in PBS (***figure 4***).

However, in aggregate these data demonstrate the feasibility of detecting CTCs *in vivo* by direct labeling with DiFC. Further implications and future steps to increase DiFC detection efficiency are discussed in more details below.

## Discussion

We recently developed DiFC, a new technique for detecting and enumerating very rare fluorescently-labeled circulating cells in the bloodstream in small animals. Until now, we have only used DiFC with cells that were “pre-labeled” before introduction into to the circulatory system, either by using cell lines genetically modified to express GFP or by labeling with a membrane or cytosol dye [32–33]. While useful for many mouse models of cancer metastasis, this ultimately restricts the use of DiFC to cultured cell lines. Therefore, we are interested in developing a receptor-based fluorescence labeling approach [47], since this could greatly extend the preclinical utility of DiFC. In addition, because DiFC is inherently scalable to larger limbs and tissues in combination with FDA approved fluorescent probes, this could ultimately open the possibility of use of DiFC in humans for enumeration of CTCs.

To achieve this, there were two main considerations for use of a FR targeted molecular probes. First, because blood is a complex mixture of many cell types the specificity and affinity for CTCs is critical. As shown in ***figures 2 and 3***, both EC-17 and Cy5-PEG-FR demonstrated extremely bright labeling in simple cell culture. However, in whole blood EC-17 performed significantly better than Cy5-PEG-FR, exhibiting was substantial separation in fluorescence labeling between FR+ CTCs and non-target cells (***figure 4***). This is consistent with previously published work with small molecular weight folate receptor targeted probes [48]. In contrast, the larger molecular weight Cy5-PEG-FR experienced significant non-specific uptake in whole blood, likely by macrophages (***figure 4***) making it infeasible as an *in vivo* injectable molecular probe based on red-fluorescence alone.

The second major consideration was the general issue of the detectability of fluorescence from a single-cell. As noted, DiFC works with diffuse photons in relatively deeply-seated (1-2 mm depth) large blood vessels. We previously estimated that DiFC requires labeling with approximately 10^5^ fluorescent molecules per cell for detectability [22]. It is well established that blue light experiences significantly more attenuation and scatter in biological tissue than red light in general, and in the specific case of DiFC (1-2 mm deep) can result in loss of approximately 50% sensitivity [33]. As such, it was unclear if EC-17 receptor-labeled CTCs would be sufficiently bright for detection with DiFC, as opposed to GFP-expressing CTCs which are generally very bright [22].

The mouse experiments performed here demonstrated that cells that were labeled in culture prior to injection were readily detectable by DiFC with high SNRs. When the cells and probe were injected separately (“*in vivo* labeling”), we were able to detect several cells, but at a count rate of approximately 12.5% compared to pre-labeled cells. This implies that many cells were labeled below the sensitivity level of DiFC, and as we noted could be explained by a reduction in EC-17 uptake by a factor of 5-6 compared to in cell culture. We tried doubling the injected quantity of EC-17 (to 50 μg) but this simply increased the background signal unacceptably, and yielded no appreciable increase in peak SNR.

Nevertheless, in combination these data provided proof-of-concept for use of FR targeted probes for DiFC. We next plan to pursue DiFC instrument and signal processing improvements to improve detection sensitivity. Moreover, a red or NIR small molecular version of EC-17 including OTL-38 may improve detectability due to more favorable tissue optics compared to blue light. This work also opens the use of DiFC to other molecular probes that target alternate cell-surface receptors. Finally, because a number of FR receptor targeted probes are in advanced clinical trials, this opens the exciting possibility of ultimately using DiFC in humans to enumerate CTCs directly in the bloodstream.

## Acknowledgements

The authors thank Dr. Xuefei Tan and Mr. Peter Bartosik for their assistance in performing some DiFC experiments.

## Availability of Materials and Data

The datasets generated and analyzed during the current study, as well as the analysis code (written in Matlab) are available from the corresponding author on reasonable request.

## References

1. Steeg PS, Theodorescu D (2008) Metastasis: a therapeutic target for cancer. Nat Clin Pract Oncol 5:206–219.

2. Gupta GP, Massague J (2006) Cancer metastasis: building a framework. Cell 127:679–695.

3. Pantel K, Speicher MR (2016) The biology of circulating tumor cells. Oncogene 35:1216–1224.

4. Alix-Panabieres C, Pantel K (2013) Circulating tumor cells: liquid biopsy of cancer. Clin Chem 59:110–118.

5. de Bono JS, Scher HI, Montgomery RB, et al. (2008) Circulating tumor cells predict survival benefit from treatment in metastatic castration-resistant prostate cancer. Clin Cancer Res 14:6302–6309.

6. Hayes DF, Cristofanilli M, Budd GT, et al. (2006) Circulating tumor cells at each follow-up time point during therapy of metastatic breast cancer patients predict progression-free and overall survival. Clin Cancer Res 12:4218–4224.

7. Liu MC, Shields PG, Warren RD, et al. (2009) Circulating tumor cells: a useful predictor of treatment efficacy in metastatic breast cancer. J Clin Oncol 27:5153–5159.

8. Cohen SJ, Punt CJ, Iannotti N, et al. (2008) Relationship of circulating tumor cells to tumor response, progression-free survival, and overall survival in patients with metastatic colorectal cancer. J Clin Oncol 26:3213–3221.

9. Toss A, Mu Z, Fernandez S, Cristofanilli M (2014) CTC enumeration and characterization: moving toward personalized medicine. Ann Transl Med 2:108.

10. Allan AL, Keeney M (2010) Circulating tumor cell analysis: technical and statistical considerations for application to the clinic. J Oncol 2010:426218.

11. Mader S, Pantel K (2017) Liquid Biopsy: Current Status and Future Perspectives. Oncol Res Treat 40:404–408.

12. Shields CWt, Reyes CD, Lopez GP (2015) Microfluidic cell sorting: a review of the advances in the separation of cells from debulking to rare cell isolation. Lab Chip 15:1230–1249.

13. Lalmahomed ZS, Kraan J, Gratama JW, Mostert B, Sleijfer S, Verhoef C (2010) Circulating tumor cells and sample size: the more, the better. J Clin Oncol 28:e288–289; author reply e290.

14. Tibbe AG, Miller MC, Terstappen LW (2007) Statistical considerations for enumeration of circulating tumor cells. Cytometry A 71:154–162.

15. Williams AL, Fitzgerald J, Niedre M (2019) Short-term circulating tumor cell dynamics in mouse xenograft models and implications for liquid biopsy. bioRxiv.

16. Wong KH, Sandlin RD, Carey TR, et al. (2016) The Role of Physical Stabilization in Whole Blood Preservation. Sci Rep 6:21023.

17. Wagar EA, Stankovic AK, Raab S, Nakhleh RE, Walsh MK (2008) Specimen labeling errors: a Q-probes analysis of 147 clinical laboratories. Arch Pathol Lab Med 132:1617–1622.

18. Delahaye M, Lawrence K, Ward SJ, Hoare M (2015) An ultra scale-down analysis of the recovery by dead-end centrifugation of human cells for therapy. Biotechnol Bioeng 112:997–1011.

19. Azab AK, Hu J, Quang P, et al. (2012) Hypoxia promotes dissemination of multiple myeloma through acquisition of epithelial to mesenchymal transition-like features. Blood 119:5782–5794.

20. Tuchin VV, Tarnok A, Zharov VP (2011) In vivo flow cytometry: a horizon of opportunities. Cytometry A 79:737–745.

21. Georgakoudi I, Solban N, Novak J, et al. (2004) In vivo flow cytometry: a new method for enumerating circulating cancer cells. Cancer Res 64:5044–5047.

22. Hartmann C, Patil R, Lin CP, Niedre M (2017) Fluorescence detection, enumeration and characterization of single circulating cells in vivo: technology, applications and future prospects. Phys Med Biol 63:01TR01.

23. Suo Y, Gu Z, Wei X (2019) Advances of in vivo Flow Cytometry on Cancer Studies. Cytometry A.

24. He W, Wang H, Hartmann LC, Cheng JX, Low PS (2007) In vivo quantitation of rare circulating tumor cells by multiphoton intravital flow cytometry. Proc Natl Acad Sci U S A 104:11760–11765.

25. Novak J, Georgakoudi I, Wei X, Prossin A, Lin CP (2004) In vivo flow cytometer for real-time detection and quantification of circulating cells. Opt Lett 29:77–79.

26. Zhong CF, Tkaczyk ER, Thomas T, et al. (2008) Quantitative two-photon flow cytometry--in vitro and in vivo. J Biomed Opt 13:034008.

27. Fan ZC, Yan J, Liu GD, et al. (2012) Real-time monitoring of rare circulating hepatocellular carcinoma cells in an orthotopic model by in vivo flow cytometry assesses resection on metastasis. Cancer Res 72:2683–2691.

28. Zharov VP, Galanzha EI, Shashkov EV, Kim JW, Khlebtsov NG, Tuchin VV (2007) Photoacoustic flow cytometry: principle and application for real-time detection of circulating single nanoparticles, pathogens, and contrast dyes in vivo. J Biomed Opt 12:051503.

29. He Y, Wang L, Shi J, et al. (2016) In vivo label-free photoacoustic flow cytography and on-the-spot laser killing of single circulating melanoma cells. Sci Rep 6:39616.

30. Galanzha EI, Menyaev YA, Yadem AC, et al. (2019) In vivo liquid biopsy using Cytophone platform for photoacoustic detection of circulating tumor cells in patients with melanoma. Sci Transl Med 11.

31. Zettergren E, Vickers D, Runnels J, Murthy SK, Lin CP, Niedre M (2012) Instrument for fluorescence sensing of circulating cells with diffuse light in mice in vivo. J Biomed Opt 17:037001.

32. Tan X, Patil R, Bartosik P, Runnels JM, Lin CP, Niedre M (2019) In vivo flow cytometry of extremely rare circulating cells. Sci Rep 9:3366.

33. Patil R, Tan X, Bartosik P, et al. (2019) Fluorescence monitoring of rare circulating tumor cell and cluster dissemination in a multiple myeloma xenograft model in vivo. J Biomed Opt 24:1–11.

34. Pera V, Tan X, Runnels J, Sardesai N, Lin CP, Niedre M (2017) Diffuse fluorescence fiber probe for in vivo detection of circulating cells. J Biomed Opt 22:37004.

35. Leamon CP (2008) Folate-targeted drug strategies for the treatment of cancer. Curr Opin Investig Drugs 9:1277–1286.

36. Parker N, Turk MJ, Westrick E, Lewis JD, Low PS, Leamon CP (2005) Folate receptor expression in carcinomas and normal tissues determined by a quantitative radioligand binding assay. Anal Biochem 338:284–293.

37. Vlahov IR, Leamon CP (2012) Engineering folate-drug conjugates to target cancer: from chemistry to clinic. Bioconjug Chem 23:1357–1369.

38. Weitman SD, Lark RH, Coney LR, et al. (1992) Distribution of the folate receptor GP38 in normal and malignant cell lines and tissues. Cancer Res 52:3396–3401.

39. van Dam GM, Themelis G, Crane LM, et al. (2011) Intraoperative tumor-specific fluorescence imaging in ovarian cancer by folate receptor-alpha targeting: first in-human results. Nat Med 17:1315–1319.

40. O’Shannessy DJ, Somers EB, Smale R, Fu YS (2013) Expression of folate receptor-alpha (FRA) in gynecologic malignancies and its relationship to the tumor type. Int J Gynecol Pathol 32:258–268.

41. Vergote IB, Marth C, Coleman RL (2015) Role of the folate receptor in ovarian cancer treatment: evidence, mechanism, and clinical implications. Cancer Metastasis Rev 34:41–52.

42. Toffoli G, Cernigoi C, Russo A, Gallo A, Bagnoli M, Boiocchi M (1997) Overexpression of folate binding protein in ovarian cancers. Int J Cancer 74:193–198.

43. Tummers QR, Hoogstins CE, Gaarenstroom KN, et al. (2016) Intraoperative imaging of folate receptor alpha positive ovarian and breast cancer using the tumor specific agent EC17. Oncotarget 7:32144–32155.

44. O’Shannessy DJ, Somers EB, Maltzman J, Smale R, Fu YS (2012) Folate receptor alpha (FRA) expression in breast cancer: identification of a new molecular subtype and association with triple negative disease. Springerplus 1:22.

45. Shi H, Guo J, Li C, Wang Z (2015) A current review of folate receptor alpha as a potential tumor target in non-small-cell lung cancer. Drug Des Devel Ther 9:4989–4996.

46. O’Shannessy DJ, Yu G, Smale R, et al. (2012) Folate receptor alpha expression in lung cancer: diagnostic and prognostic significance. Oncotarget 3:414–425.

47. Pitsillides CM, Runnels JM, Spencer JA, Zhi L, Wu MX, Lin CP (2011) Cell labeling approaches for fluorescence-based in vivo flow cytometry. Cytometry A 79:758–765.

48. He W, Kularatne SA, Kalli KR, et al. (2008) Quantitation of circulating tumor cells in blood samples from ovarian and prostate cancer patients using tumor-specific fluorescent ligands. Int J Cancer 123:1968–1973.

49. Jacques SL (2013) Optical properties of biological tissues: a review. Phys Med Biol 58:R37–61.

50. Predina JD, Newton AD, Keating J, et al. (2018) A Phase I Clinical Trial of Targeted Intraoperative Molecular Imaging for Pulmonary Adenocarcinomas. Ann Thorac Surg 105:901–908.

51. De Jesus E, Keating JJ, Kularatne SA, et al. (2015) Comparison of Folate Receptor Targeted Optical Contrast Agents for Intraoperative Molecular Imaging. Int J Mol Imaging 2015:469047.

